# Accuracy in wrist-worn, sensor-based measurements of heart rate and energy expenditure in a diverse cohort

**DOI:** 10.1101/094862

**Authors:** Anna Shcherbina, C. Mikael Mattsson, Daryl Waggott, Heidi Salisbury, Jeffrey W. Christle, Trevor Hastie, Matthew T. Wheeler, Euan A. Ashley

**Affiliations:** Division of Cardiovascular Medicine, Department of Medicine, Stanford University,Stanford, CA USA; Åstrand Laboratory of Work Physiology, The Swedish School of Sport and Health Sciences, Stockholm, Sweden; Center for Inherited Cardiovascular Disease, Division of Cardiovascular Medicine, Stanford University Stanford, CA, USA; Department of Statistics, Stanford University, Stanford, CA USA; Department of Biomedical Data Science, Stanford University, Stanford, CA USA

**Author notes:** these authors contributed equally (Anna Shcherbina, C. Mikael Mattsson, Daryl Waggott). Correspondence to: Euan A. Ashley, Falk Cardiovascular Research Building, Stanford University, 870 Quarry Road, Stanford, California, 94305.

**Keywords:** mobile health, heart rate, energy expenditure, validation, fitness trackers, activity monitors

## Abstract

**Background:** The ability to measure activity and physiology through wrist-worn devices provides an opportunity for cardiovascular medicine. However, the accuracy of commercial devices is largely unknown.

**Objective:** To assess the accuracy of seven commercially available wrist-worn devices in estimating heart rate (HR) and energy expenditure (EE) and to propose a wearable sensor evaluation framework.

**Methods:** We evaluated the Apple Watch, Basis Peak, Fitbit Surge, Microsoft Band, Mio Alpha 2, PulseOn, and Samsung Gear S2. Participants wore devices while being simultaneously assessed with continuous telemetry and indirect calorimetry while sitting, walking, running, and cycling. Sixty volunteers (29 male, 31 female, age 38 ± 11 years) of diverse age, height, weight, skin tone, and fitness level were selected. Error in HR and EE was computed for each subject/device/activity combination.

**Results:** Devices reported the lowest error for cycling and the highest for walking. Device error was higher for males,greater body mass index, darker skin tone, and walking. Six of the devices achieved a median error for HR below 5% during cycling. No device achieved an error in EE below 20 percent. The Apple Watch achieved the lowest overall error in both HR and EE, while the Samsung Gear S2 reported the highest.

**Conclusions:** Most wrist-worn devices adequately measure HR in laboratory-based activities, but poorly estimate EE, suggesting caution in the use of EE measurements as part of health improvement programs. We propose reference standards for the validation of consumer health devices (http://precision.stanford.edu/).

**Abbreviations:**

(EE): Energy expenditure
(HR): Heart rate
(GEE): General estimating equation

## Introduction

Coronary heart disease is responsible for one in every four deaths in the United States. Few interventions are as effective as physical activity in reducing the risk of death yet, we have met limited success in programs designed to help individuals exercise more. In weight loss studies, clear benefit derives from simple documentation of caloric intake,^1^ but data are less clear on the benefit of documenting exercise time and calorie expenditure.

Microelectromechanical systems such as accelerometers and Light Emitting Diode (LED)- based physiological monitoring have been available for decades.^2–7^ More recent improvements in battery longevity and miniaturization of the processing hardware to turn raw signals in real time into interpretable data led to the commercial development of wrist worn devices for physiological monitoring. Such devices can provide data directly back to the owner and place estimates of heart rate (HR) and energy expenditure (EE) within a consumer model of health and fitness. Unlike clinically approved devices, however, validation studies are not available to practitioners whose patients commonly present acquired data in the hope that it may enhance their clinical care. Indeed, certain health care systems have developed processes to bring such data directly into the medical record.^8–10^ Thus, validation data on new devices and a forum for the ready dissemination of such data are urgent requirements.

Prior studies of wrist worn devices have focused on earlier stage devices, or have focused exclusively on HR or exclusively on estimation of energy expenditure. Some made comparisons among devices without reference to an FDA-approved gold standard. None proposed an error model or framework for device validation. In response to this need, we formulated an approach to the public dissemination of validation data for consumer devices (http://precision.stanford.edu/). The website is one answer to the challenge of rapid technological advance and algorithm/product cycle upgrades. We present here the first data from this study, derived from laboratory testing of consumer wrist worn devices from the most commercially successful manufacturers. We test devices in diverse conditions on diverse individuals, and present the data and our recommendations for error modeling.

## Methods

### Patient Involvement

Patients, service users, carers, and lay people were not involved in the design or execution of the study. Study participants were recruited from the general population via word-of-mouth and e-mail notifications.

### Devices

Following a comprehensive literature and online search, 45 manufacturers of wrist worn devices were identified. Criteria for inclusion included: wrist worn watch or band; continuous measurement of HR, stated battery life >24 hours, commercially available direct to consumer at the time of the study, one device per manufacturer. Eight devices met the criteria; Apple Watch, Basis Peak, ePulse2, Fitbit Surge, Microsoft Band, MIO Alpha 2, PulseOn, and Samsung Gear S2. Multiple ePulse2 devices had technical problems during pre-testing and were therefore excluded. All devices were bought commercially and handled according to the manufacturer’s instructions. Data were extracted according to standard procedures described in the Supplementary Materials.

Participants were tested in two phases. The first group included the Apple Watch, Basis Peak, Fitbit Surge and Microsoft Band. The second group included the MIO Alpha 2, PulseOn and Samsung Gear S2.

Healthy adult volunteers (age ≥18) were recruited for the study through advertisements within Stanford University and local amateur sports clubs. From these interested volunteers, study participants were selected to maximize demographic diversity as measured by age, height, weight, BMI, wrist circumference, and fitness level. In total, 60 participants (29 men and 31 women) performed 80 tests (40 with each batch of devices, 20 men and 20 women). Participant characteristics are presented in Table 1.

**Table 1:** Participant characteristics. Values are means (min − max). Skin tone rating by Fitzpatrick scale. VO_2_max (maximal oxygen uptake) was either measured at incremental test to exhaustion or estimated from submaximal cycling using the Åstrand nomogram.

|  | Men (n=29) | Women (n=31) |
| --- | --- | --- |
| Age (years) | 40 (21 - 64) | 37 (23 - 57) |
| Body mass (kg) | 80.1 (53.9 - 130.6) | 61.7 (47.8 - 89.2) |
| Height (cm) | 179.0 (159 - 190.0) | 165.9 (154.4 - 184.2) |
| Body mass index (kg/m<sup>2</sup>) | 24.9 (20.7 - 39.3) | 22.4 (17.2 - 28.8) |
| Skin tone (scale 1 - 6) | 3.7 (1 - 5) | 3.7 (1 - 6) |
| Wrist circumference (cm) | 17.3 (16.0 - 21.0) | 15.4 (13.5 - 17.5) |
| VO<sub>2</sub>max (ml/kg/min) | 52.8 (38.2 - 66.6) | 45.3 (31.7 - 56.5) |

Skin tone at the wrist was rated independently by two of the investigators using the Von Luschan Chromatic scale (1-36), and the average rating was then transformed to the Fitzpatrick skin tone scale (1-6).^11^ Maximal oxygen uptake (VO_2_max) was measured by incremental tests in running (n = 32) or cycling (n = 6) to volitional exhaustion, or estimated from the submaximal cycling stages (n = 22) using the Åstrand nomogram (Åstrand 1960).

The study was conducted in accordance with the principles outlined in the Declaration of Helsinki and approved by the Institutional Review Board of Stanford University. All participants provided informed consent prior to the initiation of the study.

### Protocol

Participants performed the standardized exercise protocol shown in Figure 1 in a controlled laboratory setting. Participants were wearing up to four devices simultaneously and underwent continuous 12-lead ECG monitoring and continuous clinical grade indirect calorimetry (expired gas analysis) using FDA approved equipment (Quark CPET, COSMED, Rome, Italy). After being fitted with all equipment the protocol started with the participant seated for 5 min. This led to a transition to a treadmill and walking (3.0 mph at 0.5 % incline) for 10:00 min followed by faster walking (4.0 mph at 0.5 % incline) until 15:00 min, slow running (average speed 5.7 mph at 0.5 % incline, range 4.5-6.5 mph) until 20:00 min, and faster running (average speed 6.9 mph at 0.5 % incline, range 4.8-9.0 mph) until 25:00. Thereafter there was 1 min of sitting recovery, and 2 min of rest and transition to a cycle ergometer where 5 min of low intensity cycling (average work rate 88 w, range 50-100 w) until 33:00 min was followed by more intense cycling (average work rate 160 w, range 80-225 w) until 38:00 min, and 1 min of sitting recovery concluded the protocol. Both the running and cycling stages were individualized to the participants' individual fitness levels in order to maximize range of HR and EE. The last min of each stage was used for the analysis.

**Figure 1:**
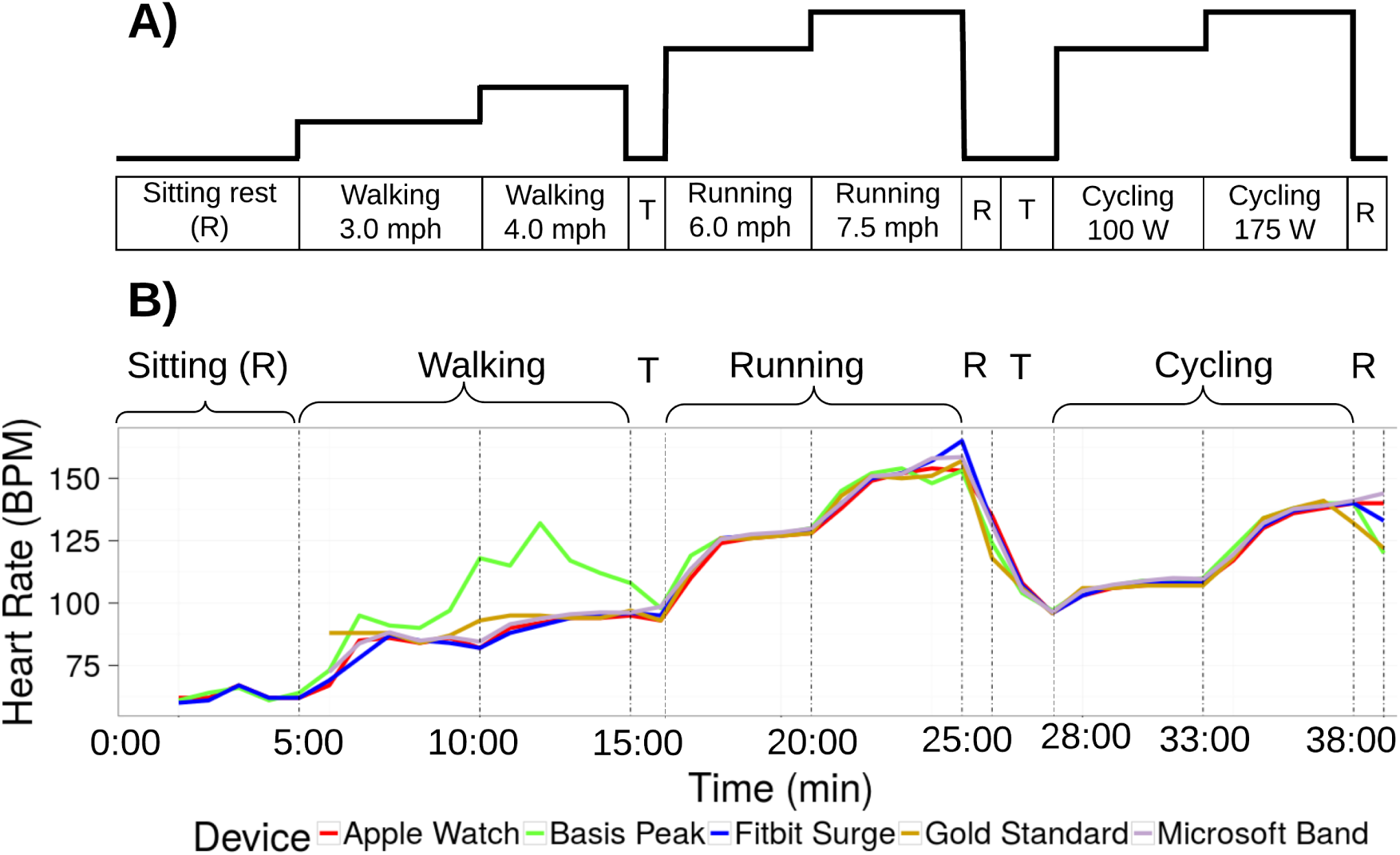
Study protocol. A) Schematic view of the protocol. Participants transition through two intensities of three modalities of exercise as shown. Walking is on a treadmill. Cycling is on a stationary bike. Activities are interspersed with brief (1 min) periods of rest “R”, and transitions between activities are indicated by “T”. B) Data from one participant wearing four devices. Data for the error analysis is derived from the last minute of each stage. Overall, error is within an acceptable range with the exception of the walking phase for one device (green line).

### Statistical analysis

Statistical analysis is described in detail in the Supplementary materials. Statistical analysis was performed separately for HR and EE. The gas analysis data from the indirect calorimetry (VO_2_ and VCO_2_) served as the gold standard measurement for calculations of EE (in kcal/min). ECG data was used as the gold standard for HR (bpm). The percent error relative to the gold standard was calculated for HR and EE using the following formula:

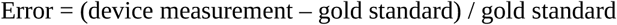

Principal component analysis, Bland-Altman analysis and various regression analyses were performed as described in the Supplementary Materials.

### Error

We determined an error rate of 5% at a p-value of 0.05 to be within acceptable limits since this approximates a widely accepted standard for statistical significance, and there is precedent within health sciences research for this level of accuracy in pedometer step counting.^12^ To gain a sense of the overall performance of each device for each parameter, a mixed effects linear regression model was utilized, allowing for repeated measurements on subjects. This was estimated using the general estimating equation (GEE) approach. First, the device type, activity type, activity intensity, and metadata confounding factors were used as inputs to a general estimating equation, with the magnitude of the error as the output variable. Second, a singular value decomposition of the dataset was performed, treating activity type/intensity as the features. Input variables were not centered, so as to find components of deviation about zero. The contribution of each feature to the first four principal components was computed to determine the degree to which it explained the variation in device measurements.

## Results

### Heart Rate (HR)

The lowest error in measuring HR was observed for the cycle ergometer task, 1.8 [0.9-2.7] *%* (all results presented as median and 95 % CI; Figure 2A), while the highest error was observed for the walking task, 5.5 [3.9-7.1] %. Six of the devices achieved a median error below 5 % for HR on the cycle ergometer task; the Samsung Gear S2 achieved a median error rate of 5.1 [2.3-7.9] %. For the walking task, three of the devices achieved a median error rate below 5 %: the Apple Watch, 2.5 [1.1-3.9] %, the PulseOn, 4.9 [1.4-8.6] %, and the Microsoft Band, 5.6 [4.9-6.3] %. The remaining four devices had median error between 6.5 and 8.8 %. Across devices and modes of activities, the Apple Watch achieved the lowest error in HR, 2.0 [1.2-2.8] %, while the Samsung Gear S2 had the highest HR error, 6.8 [4.6-9.0] % (Figure 3A and 4A).

**Figure 2.**
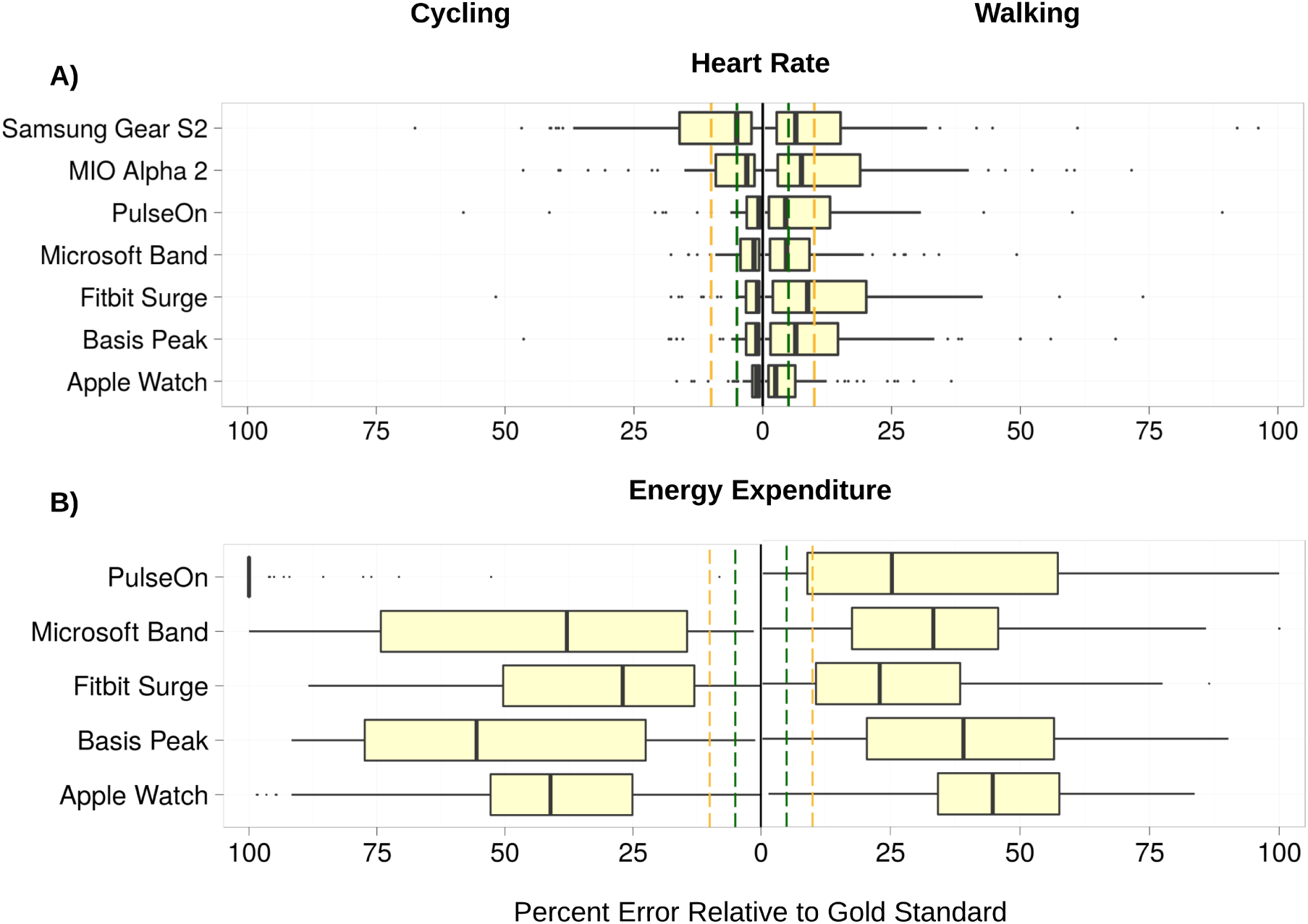
Aggregate relative error in heart rate and energy expenditure for the cycling and walking tasks -- the two tasks in the protocol with overall lowest and highest median device error, respectively. Error is calculated as *abs(Gold Standard - Device) / (Gold Standard).* The lower boundary of the boxplots indicates the 25 % quantile of data, the middle notch indicates the median data value, and the upper boundary indicates the 75 % quantile. Whiskers include all data points that fall within 1.5 IQR of the 25% and 75 % quantile values. Data points that lie further than 1.5 IQR from the upper and lower hinge values are treated as outliers, indicated by black circles. Vertical dashed green lines indicate the 5 % error threshold, while the vertical dashed yellow lines indicate the 10 % error threshold. Median heart rate error is below the 5% threshold for all but 1 devices for the cycling task, and below the 10% threshold for all devices on the walking task. Energy expenditure error rates significantly exceed the 10% threshold for all devices on both the cycling and walking tasks.

**Figure 3.**
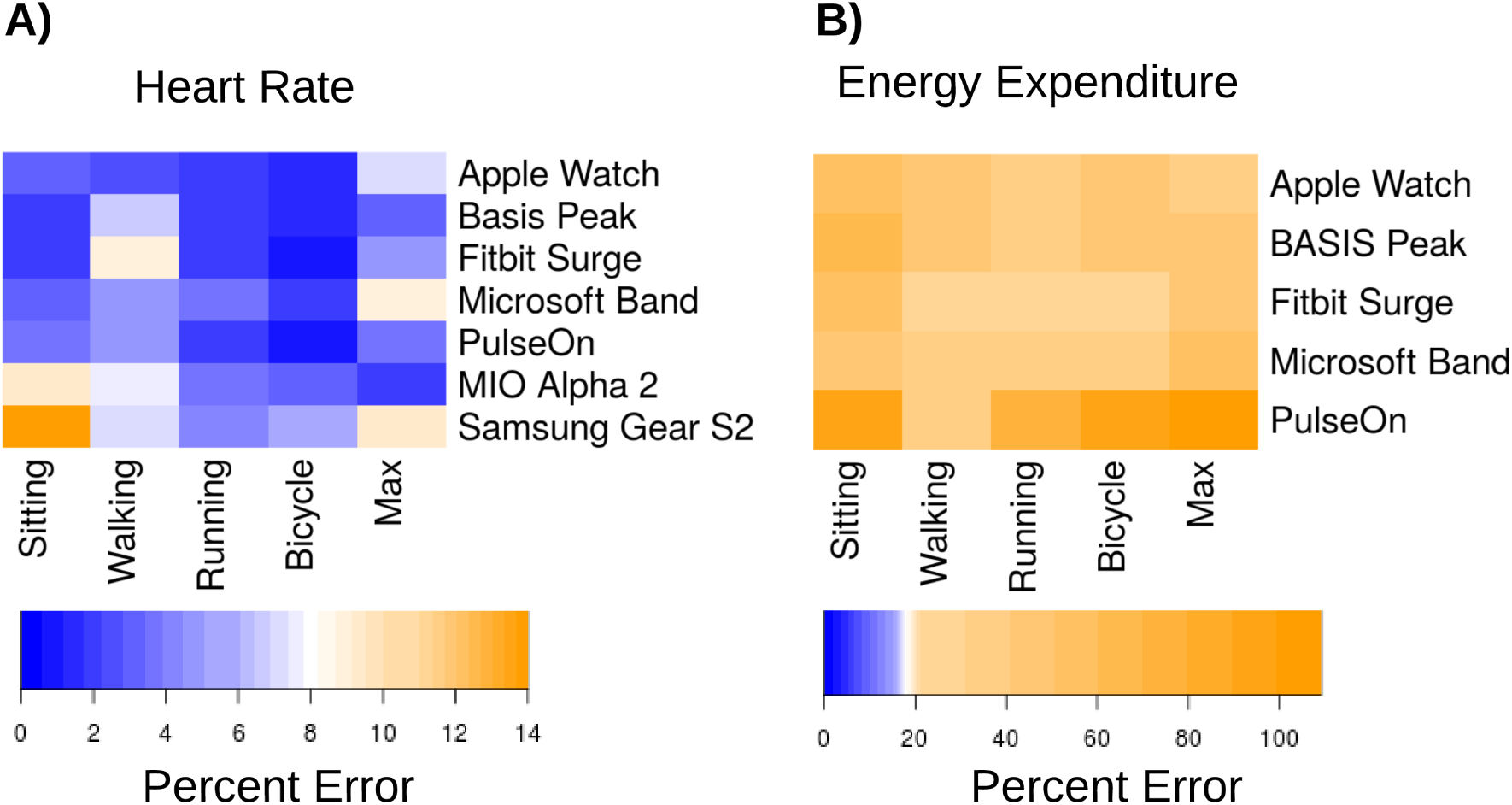
Median device error across activities. We defined an acceptable error range as <5% (dark blue). Light blue, white and yellow shading indicates error outside of this range. A) Median heart rate beats-per-minute error as a percent of the gold standard measurement. B) Median energy expenditure (kcal) error as a percent of the gold standard measurement. Note the scaling of the legend color is identical in both panels. Overall, heart rate error was within the acceptable error range for the majority of task/device combinations, but energy expenditure error exceeded the allowed threshold for all tasks and devices.

**Figure 4:**
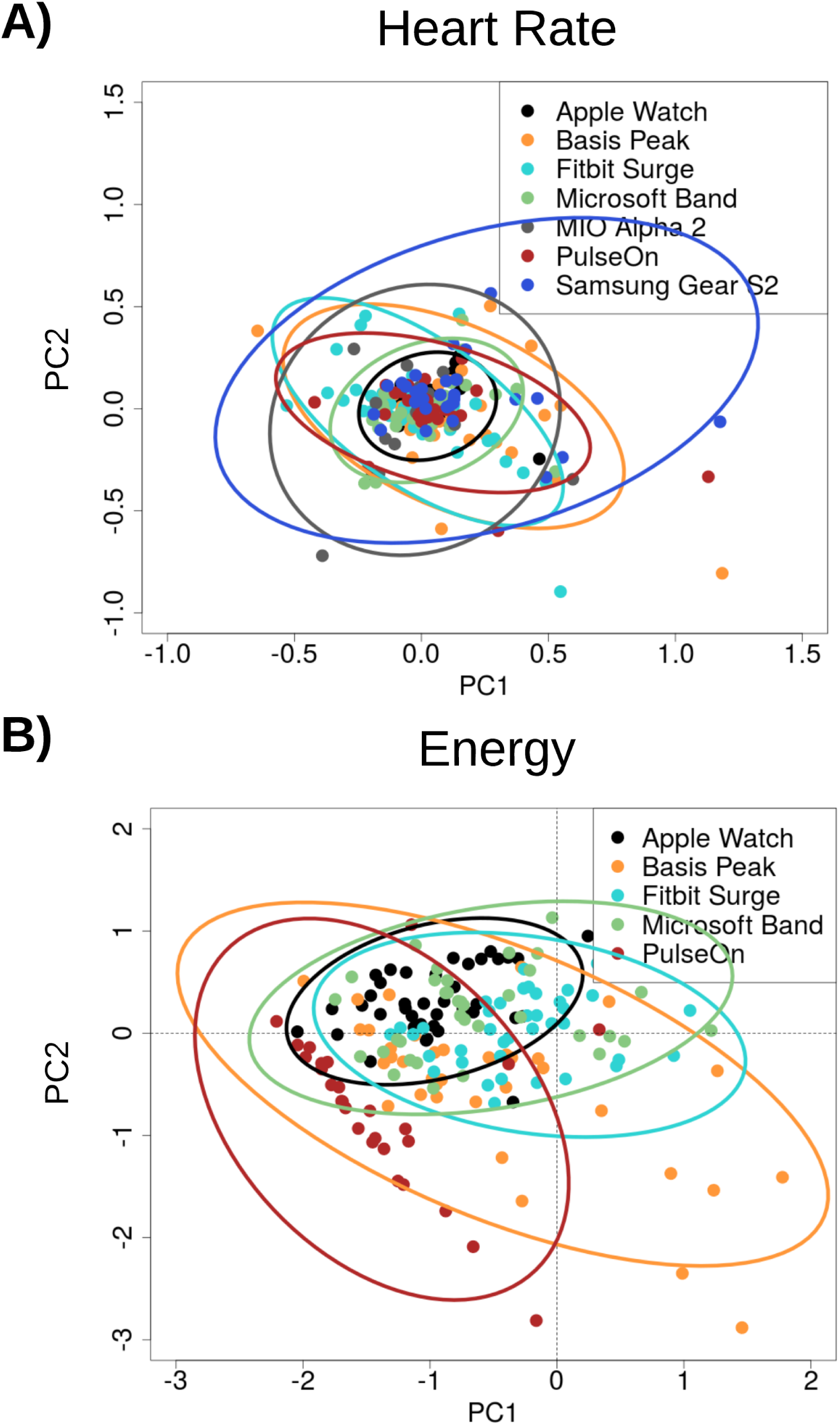
Principal component analysis of device error in (A) heart rate and (B) energy expenditure. Device errors across all activities (sitting, walking, running, cycling) were aggregated across subjects, excluding any subjects with missing data. The projection of the scaled error values on principal components 1 and 2 (PC2, PC2) are illustrated in the scatter plot, color-coded by device. Ellipses indicate the extent of the first and second principal components that encompass 95 percent of the subject error values for each device. Smaller ellipse area indicates lower variance among device error values, and data points near 0 along the PC1 and PC2 axes indicate low error. The Apple Watch had the most favorable overall error profile while the PulseOn had the least favorable.

### Energy Expenditure (EE)

Error in estimation of EE was considerably higher than for HR for all devices (Figure 2B and 3B). Median error rates across tasks varied from 27.4 [24.0-30.8] % for the Fitbit Surge to 92.6 [87.5-97.7] % for the PulseOn. For EE, the lowest relative error rates across devices were achieved for the walking (31.8 [28.6-35.0] %), and running (31.0 [28.0-34.0] %) tasks, and the highest on the sitting tasks (52.4 [48.9-57.0] %).

### Error

No evidence was found for a systematic effect of increased error for individuals across tasks or devices. Both principal component analysis and regression via the general estimating equation revealed that activity intensity and sex were significant predictors of error in the measurement of HR.. The error rate for males was significantly higher than that for females (p-value =4.56e-5, effect size = 0.044, Z=3.505) across all devices. **Supplementary Figure 1** indicates that males had on average a 4 % higher error in HR across devices and tasks. Higher VO_2_max was significantly associated with HR error on the walking task for the Microsoft Band (t=2.25) and the Basis (t=2.34). Weight, BMI, and wrist size were all negatively associated with HR error in the walking task for the Apple Watch (t=−2.53, −2.78, −2.71).

## Discussion

There are three principal findings from the current study. In a diverse group of individuals: 1) most wrist worn monitoring devices report HR with acceptable error under controlled laboratory conditions of walking, running and cycling; 2) no wrist worn monitoring devices report EE within an acceptable error range under these conditions; 3) of the devices tested, the Apple watch had the most favorable error profile while the Samsung Gear S2 had the least favorable error profile (**Supplementary Figure 3**). This study adds to the literature on wearables by including a sample of highly diverse participants, including skin tone, using FDA approved devices as a gold standard, by developing error models and by proposing a standard for clinically acceptable error.

Our finding that, HR measurements are within an acceptable error range across a range of individuals and activities is important for the consumer health environment and practitioners who might be interested to use such data in a clinical setting.

These findings are in agreement with prior work looking at fewer devices in a smaller number of less diverse individuals.^13^ In that study, HR error was within 1-9% of reference standards. In our study, six of the seven devices evaluated had a median HR error for the most stable activity, cycling, of below 5%. Covariates such as darker skin tone, larger wrist circumference, and higher BMI were found to correlate positively with increased HR error rates across multiple devices. Device error was lower for running vs walking but higher at higher levels of intensity within each modality.

In contrast with low reported error for HR measurement, no device met our prespecified error criterion for energy expenditure. This finding is also in agreement with a previous smaller study^13^ where EE estimates were up to 43% different from the reference standard. It is not immediately clear why EE estimations perform so poorly. While calculations are proprietary, traditional equations to estimate EE incorporate height, weight, and exercise modality. It is likely that some algorithms now include HR. Since height and weight are relatively fixed and HR is now accurately estimated, variability likely derives either from not incorporating heart rate in the predictive equation or from inter-individual variability in activity specific EE. There is evidence for this - for example, 10,000 steps has been observed to represent between 400 and 800 kilocalories depending on a person’s height and weight.^14^

Since devices are continually being upgraded and algorithms tuned, we created a website for sharing validation data for the community and to provide a forum for users to interact with the most up to date performance evaluations from this ongoing study (http://precision.stanford.edu/).^13,15–18^ While the FDA currently considers consumer wearable sensors such as wrist worn devices as low risk (Class 1) and therefore not subject to active regulation,^19^ they are however expected to increasingly inform clinical decision making. This makes transparency regarding benefits and limitations of paramount importance.

### Limitations

Our study has limitations. We only tested devices and algorithms that were available at the time of our study. Laboratory validation of wearable devices is a logical first step toward determining whether commercial wearables have potential use for medical applications. However the true potential of such wearables lies in their ability to provide continuous real-time monitoring outside of the clinic. This will be the focus of future research.

## Conclusions

We assessed in a controlled laboratory setting the reliability of seven wrist worn devices in a diverse group of individuals performing walking, running and cycling at low and high intensity. We found that in most settings, heart rate measurements were within acceptable error range (5%). In contrast, none of the devices provided estimates of energy expenditure that were within an acceptable range in any setting. Individuals and practitioners should be aware of the strengths and limitations of consumer devices that measure heart rate and estimate energy expenditure. We encourage transparency from device companies and consistent release of validation data to facilitate the integration of such data into clinical care. We provide a forum for the community to share such data freely to help achieve this end.

